# How to deal with internal fragment ions?

**DOI:** 10.1101/2024.04.18.589916

**Authors:** Arthur Grimaud, Maša Babović, Frederik Haugaard Holck, Ole N. Jensen, Veit Schwämmle

## Abstract

Tandem mass spectrometry of peptides and proteins generates mass spectra of their gas-phase fragmentation product ions, including N-terminal, C-terminal, and internal fragment ions. Whereas N- and C-terminal ions are routinely assigned and identified using computational methods, internal fragment ions are often difficult to annotate correctly. They become particularly relevant for long peptides and full proteoforms where the peptide backbone is more likely to be fragmented multiple times. Internal fragment ions potentially offer tremendous information regarding amino acid sequences and positions of post-translational modifications of peptides and intact proteins. However, their practical application is challenged by the vast number of theoretical internal fragments that exist for long amino acid sequences, leading to a high risk of false-positive annotations. We analyze the mass spectral contributions of internal fragment ions in spectra from middle-down and top-down experiments and introduce a novel graph-based annotation approach designed to manage the complexity of internal fragments. Our graph-based representation allows us to compare multiple candidate proteoforms in a single graph, and to assess different candidate annotations in a fragment ion spectrum. We demonstrate cases from middle-down and top-down data where internal ions enhance amino acid sequence coverage of polypeptides and proteins and accurate localization of post-translational modifications. We conclude that our graph-based method provides a general approach to process complex tandem mass spectra, enhance annotation of internal fragment ions, and improve proteoform sequencing and characterization by mass spectrometry.

## 2 Introduction

The regulation and fine-tuning of the majority of cellular processes are made possible by the vast diversity of proteoforms originating from mechanisms such as RNA splicing, post-translational modification (PTM) or protein truncation [1, 2]. The characterization of proteins at the proteoforms level is essential for the comprehensive understanding of complex cellular processes and regulatory mechanisms. While some variation of the proteins’ canonical sequence can be measured at the RNA level, the true complexity of proteoforms can only be approached by their direct characterization. Antibody-based approaches can typically only monitor single or few cooccurring PTMs, and PTM-specific antibodies have often low specificity and showcase bleedthrough [3]. For this reason mass spectrometry (MS) constitutes the main approach for proteoform characterization. More specifically, approaches that allow measuring long peptides or entire proteins such as middle-down and top-down mass spectrometry have gained a lot of interest for their capacity to identify distant co-existing PTMs [4, 5, 6].

Middle-down and particularly top-down MS come with the cost of both increased spectral complexity and expanded search space due to the length of the analytes and many possible modification sites [7, 1, 8]. Tandem mass spectra (MS2) from such approaches often result in incomplete fragment coverage of the sequence. The absence of lack of information at the MS2 level can render the determination of the sequence and the localization of PTMs ambiguous or impossible. Several methods have been employed to alleviate the difficulties of identifying large molecules in these MS approaches. Top-down or middle-down MS experiments are sometimes integrated with data from bottom-up experiments to capitalize on both approaches’ advantages [9, 10, 11]. Insufficient sequence information in top-down approaches has also been pondered, either by integrating multiple MS2 fragmentation techniques to enhance sequence coverage [12] or by performing multiple levels of fragmentation of a protein ion, known as MSn [13, 14]. A drawback of these methods is that they involve several rounds of analysis to be carried out and therefore require larger sample amounts.

The identification of peptidoform and proteoform species in middle-down and top-down MS analysis of complex samples can be compromised from the incomplete sequence coverage by fragment ions in MS2 scans. Additionally, the presence of multiple co-eluting and isomeric (or near-isobaric) species in a MS2 spectrum can lead to the absence of site-determining terminal ions therefore hindering the confident localization of PTMs [15]. While some of the issues encountered in PTM localization are caused by incomplete fragmentation of the peptide or protein backbone, it is in some cases even impossible to localize PTMs in certain chimeric spectra because of the incompleteness of information carried by terminal fragment ions.

Internal fragments would theoretically solve many of the above-mentioned issues. Internal fragments are types of fragments that occur through secondary or higher fragmentation events of the backbone. The first fragmentation of a peptide or protein yields two terminal fragments (N and C terminal). If these fragments undergo a second fragmentation event, the result is an internal fragment that contains neither termini of the original sequence. Internal fragment ions can carry important information as they can increase the coverage of the central part of the peptide or protein, where terminal fragments are usually less observed in middle-down and top-down MS. Moreover, internal fragments might “frame” specific modification sites in the sequence which can greatly improve the confidence of PTM localization.

Internal ions have been described and used early on to sequence peptides and proteins [16, 17] and for PTM identification [18, 19]. Although such product ions have been introduced, internal ions have mainly remained absent from standard proteomics workflows. With the advance of top-down proteomics, we can observe a regained interest in internal fragment utilization. Several computational approaches that emphasize curating internal ions annotation have been published and show that internal ions can be used to improve the coverage of the sequence [20, 21, 22, 23]. However, a thorough analysis of their potential has been lacking.

The difficulty in using internal fragment information resides in their annotation in the spectrum. For a given sequence of length *n* there exists *n*2-2* different positions for terminal ions but *n*(n+1)/2* positions for internal ions and many of the possible internal ions can be expected to not be generated or be below the detection threshold. Furthermore, the number of theoretical fragments multiplies when several fragment types can be attributed from a fragmentation method, as is the case in electron-transfer dissociation (ETD). Considering theoretical fragments in the annotation therefore introduces a high risk of false-positive annotation, from both annotating noise peaks or from the high number of overlapping fragments with identical or near-identical mass, and thus would disqualify the usage of internal ions.

In this work, we initially assess the challenges that arise when annotating internal ions by analyzing theoretical spectra exhibiting internal ions. We focus here on MS2 scans from ETD as it is the fragmentation method commonly used in sequencing and PTM characterization of long polypeptides and intact proteins [24, 25]. As the sequence length is the main determining factor for the complexity of the set of internal fragment ions, we provide a comparison of challenges for bottom-up, middle-down, and top-down approaches. Accounting for these challenges, we present a graph-based method to represent and annotate all fragments in non-deconvoluted spectra of long species. We present annotation results using our method first on experimental middle-down spectra and then apply it to top-down spectra. By focussing on histone proteins, we tackle the challenges arising from heavily modified proteoforms. The fragment annotation from the approach presented allows us to make use of complementarity between terminal and internal ions, to show that internal ions are indeed generated when the fragmentation energy (or reaction time) is sufficient. We also propose a method to determine optimal fragmentation parameters for the generation of internal and terminal ions. Finally, the annotation results from the developed method were used to statistically assess how beneficial internal fragments are to PTM localization and proteoform characterization.

## 3 Experimental Procedures

### Middle-down: Histone H3.1 n-term tail

Unmodified N-terminal histone H3.1 tail peptides (residues 1-50) were synthesized by F-Moc chemistry as previously described [26]. The peptide standard was directly infused at a concentration of 2 pmol/µL in 0.1% formic acid using a Borosilicate Emitters (ThermoFisher, USA, ref: ES387) by applying positive air pressure via a syringe at roughly 1µL/min. Peptides were infused into an OrbitrapTM FusionTM LumosTM (Thermo) with a spray voltage of 2200V and ion transfer tube temperature of 275 °C. MS1 spectra were obtained in the Orbitrap at 120.000 resolution, AGC target of 1E6, max ion injection time of 50ms, and 2 averaged microscans. MS/MS spectra were obtained in the Orbitrap with a mass resolution of 30.000, AGC target of 1E5, max ion injection time of 200ms, and 3 microscans per MS/MS event. Additional fragmentation parameters such as ETD reaction time, HCD fragmentation energy, and supplemental activation energy were varied to assess fragmentation patterns.

### Top-down: Histone H3.1 & H4, Filgrastim

Histone H3.1 (H2292, Sigma) and histone H4 (H2667, Sigma) were desalted using an in-house packed pipette tip with Poros R1 reversed-phase material (Thermo Fischer, USA). Desalted proteins were diluted to 2.5 µM in water: formic acid (50:50:0.5) solvent and directly infused into Orbitrap Eclipse mass spectrometer with HESI source (Thermo Fischer, USA) at 2 µL/min flow rate, with cone voltage 3500V, and ion transfer tube temperature 300 C*^◦^*. Targeted MS2 methods were created using topdownr version 1.15.1 and XmlMethodChanger [27]. For each protein, 108 unique fragmentation settings were used. For the filgrastim analysis protocol refer to Babovic et al [12]. The specific fragmentation method and parameters of MSMS spectra in the analysis are specified in the figure captions.

### Generation of theoretical fragment lists

Theoretical fragments lists were generated for peptides of different lengths using a custom R script. For a better representation of real datasets, peptides of length ranging from 8 to 25 amino acids were sampled at random from the human tryptic peptide list obtained from PeptideMapper’s benchmark dataset at (https://github.com/compomics/PeptideMapper-Benchmark/tree/master/peptide-lists) [28]. For longer peptides, with lengths spanning from 30 to 200 amino acids, random sampling was performed from the human reference proteome (UP000005640, August 2023).

### Spectra deconvolution and merging

Raw spectra were centroided and converted to mzMl format using ThermoRawFileParser (v1.7.3) [29]. When specified, spectra deconvolution (deisotoping and charge deconvolution) has been performed using Xtract via Proteome Discoverer (v2.5.0.400) (Thermo Scientific). The method used to combine multiple spectra in a consensus spectrum is a Python re-implementation of the “combine spectra” function of the Msnbase R package [30]. Peaks of multiple spectra are combined if their mass-to-charge ratio (m/z) difference is smaller than the specified tolerance. For each dataset, the tolerance was determined from the scattering of m/z differences as shown in the m/z calibration histogram in Supplementary Figure S3. The mean of the intensity and the mean of m/z were used to compute the peaks of the consensus spectra. Filtering was applied to remove peaks that appear in less than 10% of the spectra used in the merging.

### Graph-based annotation of spectra

We implemented a hierarchical structure where the parent node(s) represents the precursor ions and stores information about the candidate peptidoforms (or proteoforms). Branching from that precursor node are position nodes that represent all possible fragment positions for both terminals and internal fragments. From these nodes, successive nodes were created to display the fragment types, the possible charges, and the different isotopes. The fragment types were determined by the fragmentation method used. For the ETD spectra analyzed in this study zdot, z+1, z+2, c, and cdot ion caps were considered (see Suppl. Tab. S1 for mass calculations) [31, 32]. The maximum charge considered for the fragments was determined as the charge of the precursor selected from fragmentation. Isotopes of a fragment were considered until the expected abundance of that isotope fell under the detection threshold. In practice, the expected intensities of the isotopes of a fragment were calculated by using the intensity of the peak matching the most abundant isotope of that fragment and the theoretical probabilities of the isotopes. If the calculated expected intensity was inferior to the least intense peak in the experimental spectrum, the isotope was not considered. Through these different types of nodes, each path from a “position” node to an “isotope” node represents a specific fragment ion and potential peak that was then matched to the experimental spectrum (Fig. 1A). A tolerance of 10 ppm was considered for the matching.

**Figure 1:**
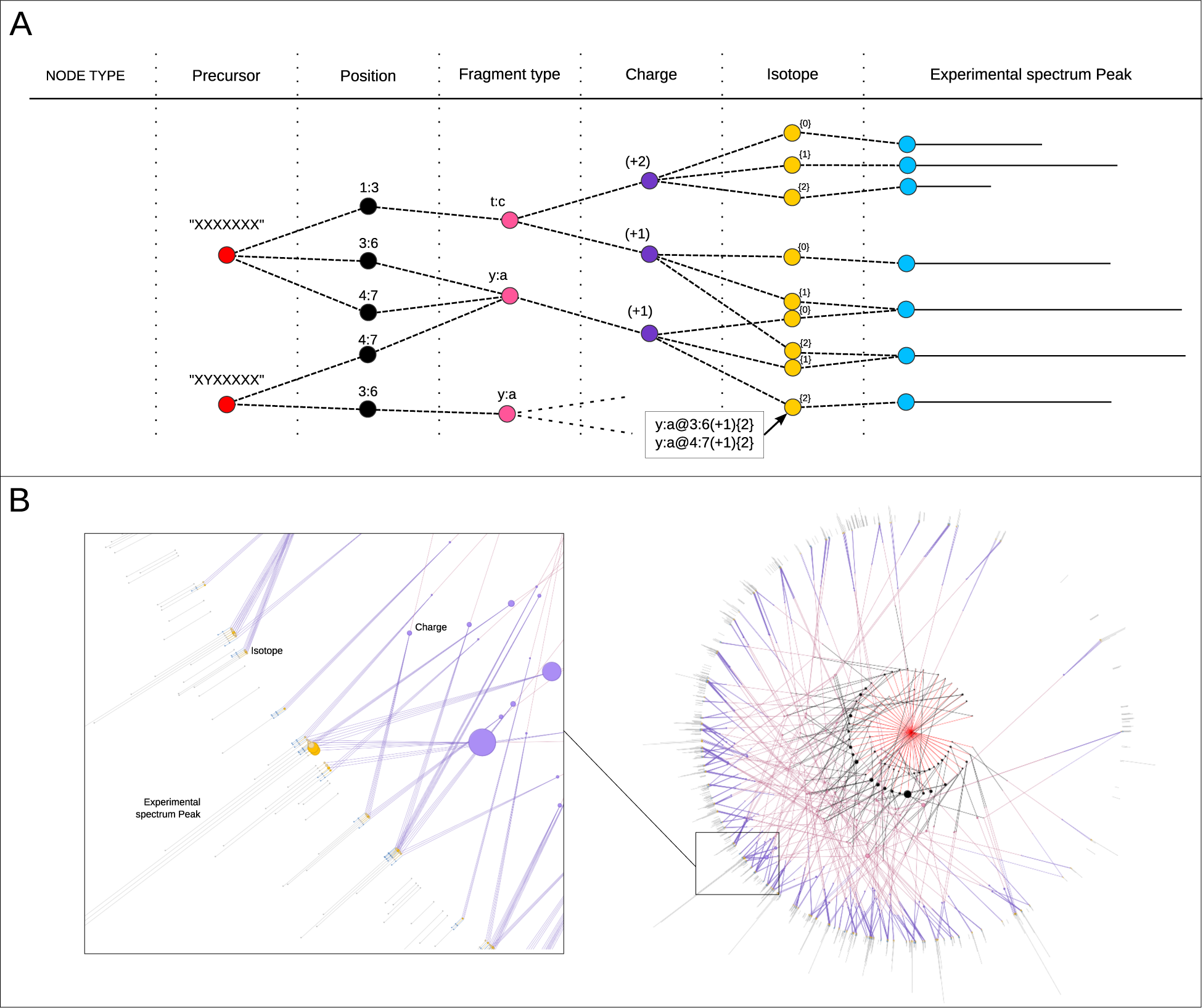
Graph-based annotation of non-deconvoluted mass spectra. The graph structure presented in panel **A** is used to represent all possible fragments of a sequence. The root node(s) correspond to the candidate peptidoform(s) or proteoform(s) and each child node corresponds to the different possible positions of fragments. From these “position” nodes “fragment type”, “charge” and “isotope “nodes represent the different combinations of fragment properties therefore representing the collection of theoretical peaks for that precursor. Given the specified mass tolerance, the isotope nodes are then matched to nodes representing the experiment. Two “fragment type” nodes from different position nodes will be merged as one if they have the same mass, this avoids overlapping redundant annotation (i.e ‘y:a’ from ‘3:6 and ‘4:6, panel **A**). This graph structure has the advantage of being able to represent theoretical fragments for multiple precursors. Identical fragments of different precursors will be merged to avoid redundant annotation. **B:** Representation of the graph for the annotation of a Histone H3 N-terminal tail spectrum (50 amino acid long), experimental peaks are displayed on the out circle, and the precursor node in the center in red.

The fragment annotations were scored by comparing the theoretical isotopic distribution to the observed using cosine similarity. Computation of theoretical isotopic distribution was performed using the BRAIN algorithm [33, 34]. In the case where multiple theoretical fragment isotopic patterns overlap, an optimization problem was solved to assign weights to each overlapping fragment in the function of their contribution to the observed isotopic pattern in the spectrum. In the case where the contribution of a fragment’s theoretical isotopic pattern did not improve the fit between the observed and theoretical isotope pattern, the weight of this fragment was set to 0 and the annotation was discarded. Once the graph has been generated, the fragments matched and poorly scoring annotations filtered out, and the intensities assigned to each “isotope” node were propagated to the parent nodes. The intensity assigned to a parent node is the sum of all its children nodes. In the case where a node has multiple parents, for instance when a “fragment-type” node is assigned to multiple positions, the intensity was distributed evenly across the parent nodes to consider the overlap in mass and the incapacity to select one fragment annotation over the other. In the case where an internal node overlaps with a terminal node, the terminal node was retained as considered more likely.

This graph representation allows the theoretical spectrum of multiple candidate peptide forms (or proteoforms) to be represented in the same graph. For that two or more precursor nodes were added to the graph and subsequent nodes representing fragments are generated from all the precursors. In the case where fragments are shared across candidate species, these nodes were “merged” to limit redundancy (Fig. 1A). The implementation of this approach made use of the Python package NetworkX (version 2.2) [35]. A diagram of steps for the entire annotation procedure in presented in supplementary figure S1

## 4 Results

We aim to address three main aspects of internal fragment annotation. Specifically, (1) What are the principal challenges in annotating internal fragment ions, and how do these challenges manifest across different MS approaches? (2) How can these challenges be mitigated with an improved annotation approach? (3) What is the value of information carried by internal fragments, and how does this enhance the ability to characterize proteoforms, particularly in the context of localizing PTMs?

### Internal fragment ions pose new challenges to spectrum annotation

The number of theoretical internal fragment ions exponentially increases with the sequence length (Fig. 2A). This poses two main challenges: firstly, the m/z range covered by theoretical fragments becomes important for long peptides, possibly leading to annotation to noise or contaminations (Fig. 2B), and secondly, an increasing number of theoretical fragments have overlapping masses, making it difficult or impossible to determine which annotation to assign. While the first issue can be addressed by increasing mass resolution, the second persists as most overlapping fragments are isomeric, and increased mass resolution may only help with distinguishing near-isobaric fragments (Suppl. Fig. S2). These issues are expected to be neglectable in spectra of short peptides from bottom-up MS experiments. However, in spectra in middle-down or top-down MS, the risk of annotation of noise and fragment overlap cannot be omitted. For this reason, we developed a graph-based annotation, that aims to reduce annotation to noise and consider fragment mass overlaps.

**Figure 2:**
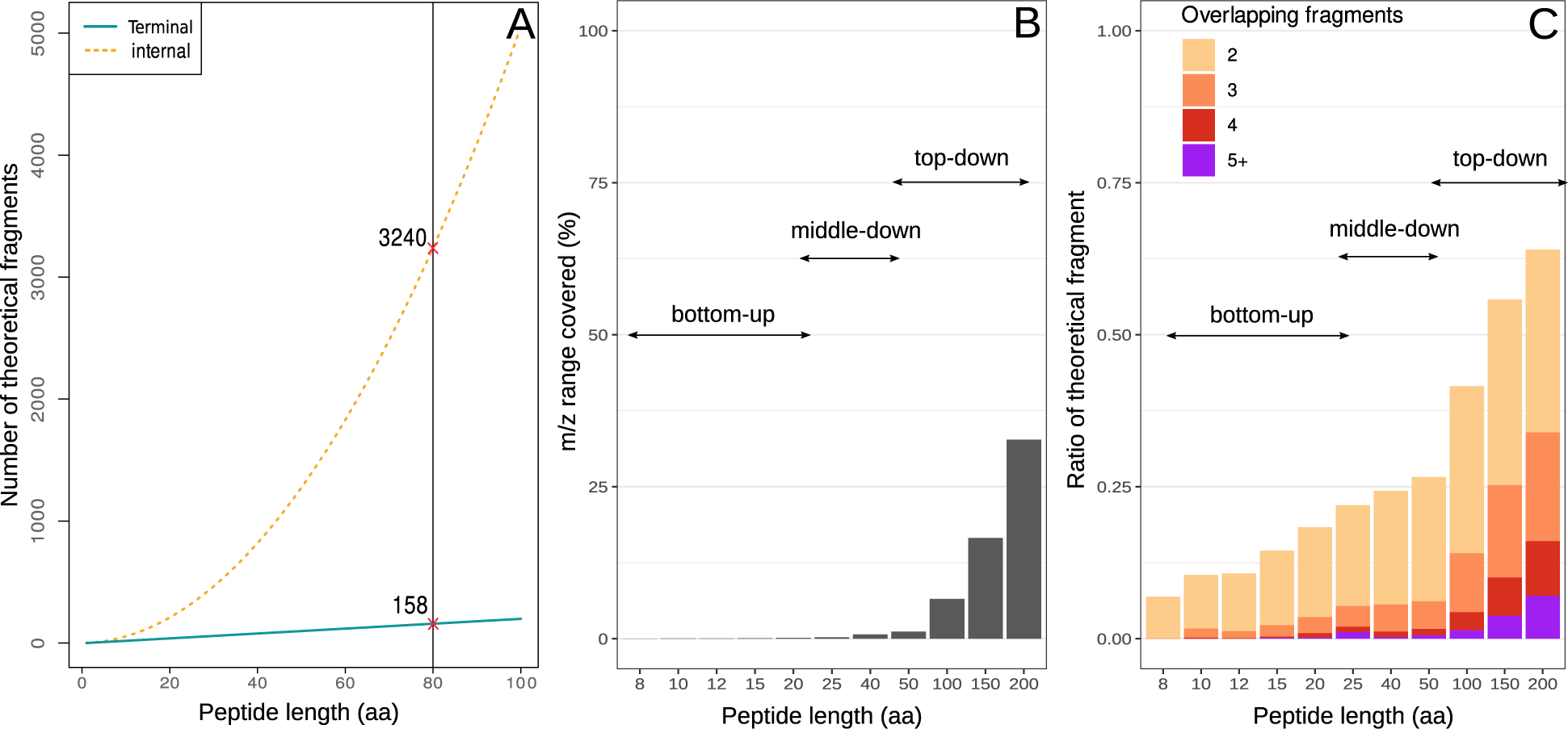
Increased sequence length drastically increases fragment annotation complexity. **A:** The number of possible internal fragments grows exponentially with the fragment length leading to a substantial portion of the m/z range of a spectrum to be “covered” by theoretical fragments. **B:** The coverage is here defined as the proportion of the m/z range of the spectrum where a theoretical fragment can be located, taking into account the specified error tolerance. Higher coverage of the m/z range indicates that more noise peaks are expected to be matching to at least one theoretical fragment. **C:** A very high number of theoretical fragments leads to many of them having overlapping masses. (Theoretical fragments for typical ETD spectra, MS2 tolerance of 5 PPM)

### Which internal ions annotation to retain and how to optimize internal fragment generation

As shown previously, adding internal ions in the annotation of spectra greatly increases the risk of false-positive annotations. Therefore, before annotating internal ions, the presence and abundance of such fragment types in the spectra should be assessed, to ponder whether enough internal ions are present to provide valuable information or whether fragmentation parameters can be tuned to increase the rate of formation of internal ions. For that, we will use the dataset consisting long histone H3 N-terminal peptides from a middle-down experiment as they allow a more precise annotation of internal ions than top-down spectra from longer proteoforms.

To assess the presence of internal ions in middle-down spectra of histones N-terminal tail, we looked into the correlation between the summed fragment intensities on the N and C terminal sides of each fragmentation site (denoted N/C correlation). We hypothesize that in the case where each peptide undergoes only one fragmentation event, the intensity in both directions of the fragmentation site should be more or less equal. Here we assume that the amount of signal lost because of neutrally charged fragments can be neglected for long peptides with high charge states. We observed that the N/C intensities are not correlated when examining terminal fragments in spectra with sufficiently high fragmentation energy (Fig. 3). This suggests that the peptides underwent more than one fragmentation event and that the intensity correlation between N and C terminal ions is lost because of the formation of internal fragments. As each of the fragment ions in the annotation was scored according to their similarity to the expected isotope distribution, we can accept internal ion annotations successively by decreasing the score threshold and then re-measuring the N and C terminal intensities considering intensities of internal ions(Fig. 3A).

**Figure 3:**
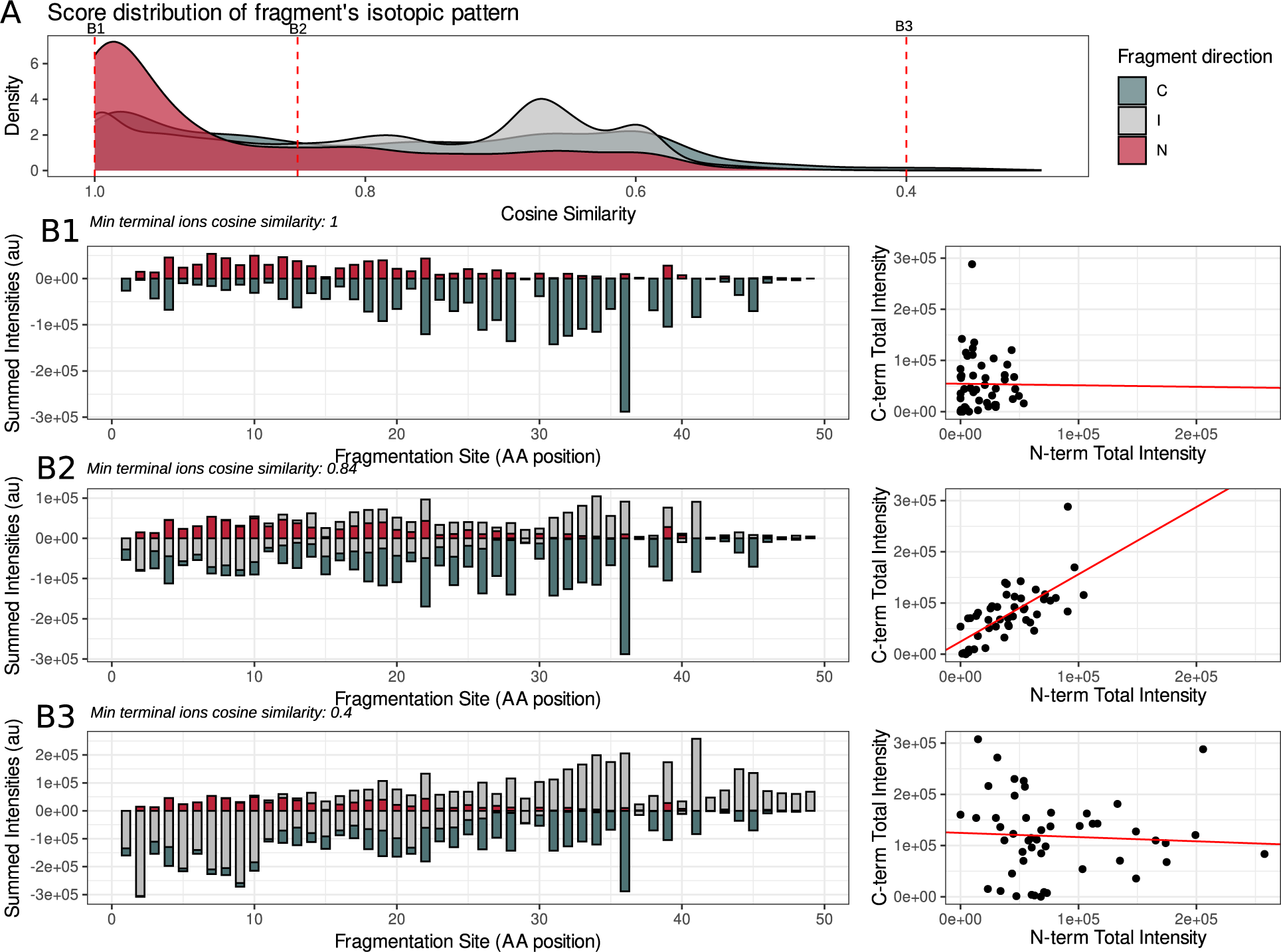
Internal ions account for discrepancies between N and C terminal fragment intensity. **A:** Annotation of the peak-picked spectrum (consensus spectrum from 45 histone H3 n-terminal tail spectra, ETD reaction time 40; supplementary activation 20) with the fragmentation graph approach enables to score each fragment annotation according to its cosine similarity score calculated between the expected and measured isotopic pattern of fragment ions. **B1:** When only terminal fragments are included, a discrepancy between N and C terminal fragment intensities is observed as short fragments are more likely to be observed with this fragmentation setting, or in other other words, N terminal fragments are more intense on the N termini of the peptide and C terminal on the C termini. Such discrepancy can be explained by the presence of internal fragment ions that are not included and account for intensity on the opposite side of the fragmentation site resulting in a lack of “N/C correlation” (Pearson coefficient of correlation. -0.01). **B2:** The balance between N and C terminal intensity of the fragmentation site is corrected as internal fragments are added by lowering the cosine similarity threshold. The correlation reaches a maximum (Pearson coefficient of correlation: 0.69), at a threshold of 0.84 before decreasing for lower score thresholds (**B3**, Pearson coefficient of correlation: -0.06). This suggests that low-scoring fragment annotations are false-positive as they do not contribute to correcting the N/C term balance.

As more internal ions are added, the N/C correlation increases until reaching its maximum (Fig. 3B1 and B2). After that, including low-scoring internal fragment ions resulted in a decrease in the N/C correlation (Fig. 3B3) suggesting that most of these low-score annotations were false positives. Through this approach, we can determine the optimal score at which the maximum N/C intensity correlation is achieved. We do not observe this increase in N/C correlation for spectra with lower fragmentation energy where one does not expect many secondary fragmentation events (Fig. S6 B).

The measurement of changes in N/C correlation can be used to determine optimal fragmentation settings yielding a maximum of internal ions resulting from two fragmentation events. Figure 4 showcases this approach in middle-down data. Unmodified N-term tails of H3 (sequence length of 50) were analyzed with ramping ETD reaction time (10ms to 40ms) and different supplementary activation (denoted low and high in Figure 4 for clarity). For higher fragmentation efficiency, we observe an increase in the maximal correlation values (Fig. 4A), reaching a correlation coefficient of 0.56 (Pearson) for ETD 30 ms reaction time. Thereafter, the correlation drops for higher ETD reaction times. We hypothesize that these higher fragmentation energy settings yield fragments resulting from three or more successive fragmentation events, which leads to a loss in complementarity between terminal and internal fragment ions (Fig. 4C). Additionally, the short fragments that are yielded from high fragmentation efficiency are less likely to be charged, and therefore to be detected. In the sample analyzed here an ETD reaction time of 30 ms appears to be the optimal parameter for the generation of internal ions. It is worth noting that no significant drop in terminal ions coverage of the peptide sequence is observed for ETD 30ms compared to ETD 10ms, implying that internal ions can be used without losing relevant information at the terminal ion level (Fig 4B)

**Figure 4:**
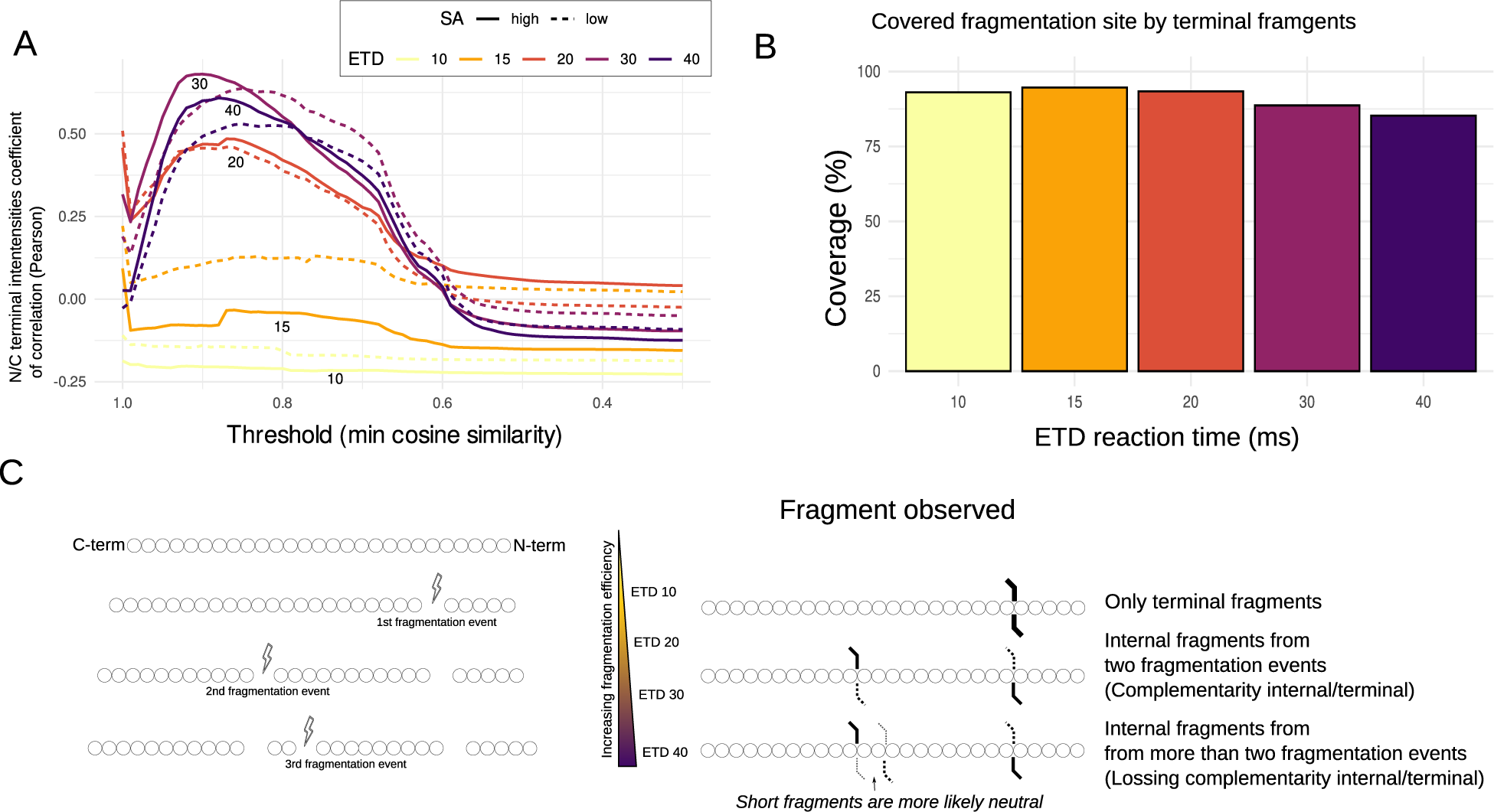
Changes of N/C intensity correlation determine optimal fragmentation settings for the generation of internal ions. **A:** The correlations between N- and C-terminal fragment intensity were compared for different internal fragment score thresholds and various fragmentation parameters fragmentation energies (colors), as well as different supplementary activation (line type). **B:** Sequence coverage by terminal fragments (average percentage of possible fragmentation sites where a terminal fragment is found). **C:** Illustration of the hypothesis behind the loss of internal/terminal fragment complementarity for higher ETD reaction times.

### In-depth annotation of spectra with a fragmentation graph approach

After showing that internal ions are indeed present within a range of fragmentation settings, we will now explore how well they can be annotated without assigning too many false positives in spectra of long peptides, to fully explore potentially powerful methods that then can be applied to very long peptidoforms and proteoforms.

The choice of a graph-based model for theoretical fragment representation and annotation is motivated by several aspects; the capacity to represent multiple layers of fragment properties in order to annotate non-deconvoluted spectra, to efficiently represent overlapping fragments, and to simultaneously include fragments from multiple peptidoforms.

We can assume that valuable spectral information is lost during the spectra deconvolution step typically performed in middle and top-down MS (Suppl. Fig. S10). Because of their low intensity, peaks in the spectra corresponding to internal fragments might be filtered out by noise removal algorithms of deconvolution tools or being confused with other ions. In addition to that the presence of hydrogen transfer products (i.e z+1, z+2, cdot) in ETD spectra [36] are yielding fragments with a ± 1 Dalton shift that can be confused for ±1 Dalton peaks from the isotopic distribution [37, 38]. Moreover, the isotope information available in non-deconvoluted spectra allows us to apply additional scoring strategies, as applied in terminal fragment annotation in TDValidator [39]. Our graph structure can efficiently represent all theoretical peaks of a given “positional” fragment by storing information about the different fragment types, charge states, and isotopes as different nodes. This representation allows to efficiently annotate fragments in non-deconvoluted spectra. The graph-based approach also takes into account the overlapping fragment annotations by allowing for spectrum peaks to be matched to multiple fragments and allows methods for sharing the intensity of such signal between multiple annotated fragments.

Another strength of this approach resides in the capacity to compute the mass of fragments at different levels of the tree, in the case where two nodes are isomeric or nearly have the same mass within the mass tolerance specified, their nodes can be easily merged into one (Fig. 1A). This improves annotation speed by avoiding redundant assignments of overlapping fragments.

Our graph representation will prove to be beneficial when the annotations of the theoretical fragments of multiple candidate species are compared within the same graph structure. Then, representing multiple candidates is beneficial to the annotation, as a considerable amount of internal fragments may overlap, which cannot be effectively accounted for in separate representations.

As demonstrated in Figure 5A, the graph-based method for annotating fragments in non-deconvoluted spectra allows representing sets of peptidoforms in a single object. The annotation of the full array of theoretical fragments within non-deconvoluted spectra significantly enhances the proportion of explained peaks and allows minimizing false-positive annotations through the filtering of fragment annotation. For instance, in the annotated spectrum of Histone 3.1 N-terminal tail shown in Figure 5B, internal fragments account for almost half of the spectral peaks and about one-quarter of the total intensity after filtering. As shown in Figure 5C and Figure S4 the annotation of internal ions drastically increases the sequence coverage. Additionally, internal fragment annotations complement terminal ion annotations by covering the central regions of the sequence, where terminal fragments are typically scarce. However, because the rate of annotated false positives is expected to be high for internal ions, additional investigation is required to determine the true added value of internal ions for proteoform characterization. Also, the visualization of annotated internal fragments in spectra from species with long sequences poses challenges due to the large amounts of fragments. Consequently, we propose and later use “fragment coverage matrices” for a clearer representation of the annotations and sequence coverage.

**Figure 5:**
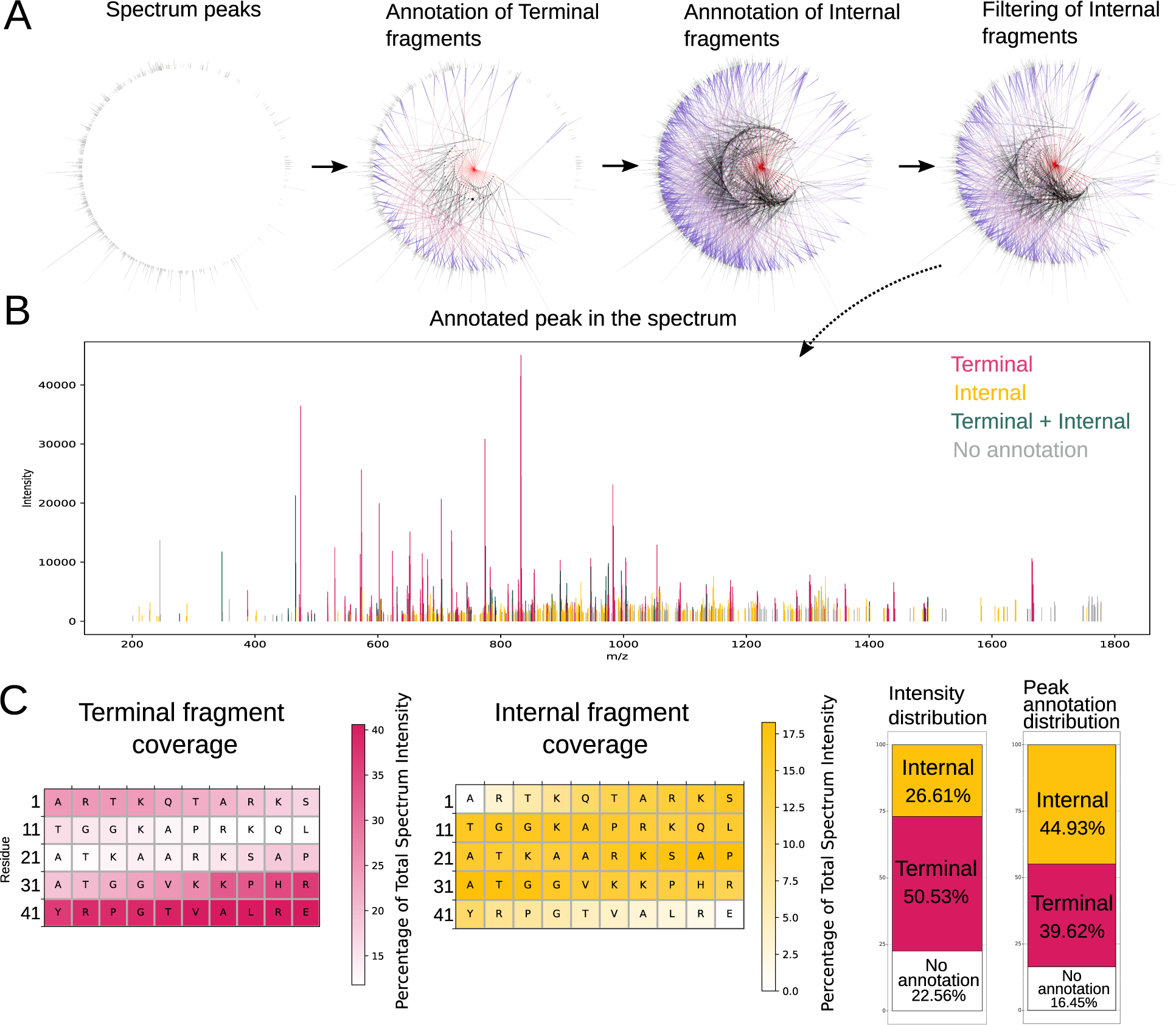
Annotation of internal fragments with the fragmentation graph approach increases information content that can be extracted from non-deconvoluted spectra. **A:** Overview of the graph for different steps of the annotation. First, the experimental peaks are represented as nodes, theoretical terminal, and internal nodes are then generated as a graph, and internal fragment annotations are filtered according to their score. **B:** The annotated spectrum illustrates the assignment of numerous internal fragments to low-intensity peaks within the spectra. **C:** The sequence coverage of terminal and internal ions is represented as the percentage of intensity in the MS2 spectrum covering each residue. The accompanying bar plots display the summed intensity and count of internal, terminal, and unassigned peaks. (Dataset: Middle-down Histone H3 n-term tail, ETD 30ms)

### Internal ions can facilitate the localization of PTMs

PTM localization is a challenging problem in complex spectra from long species, leading to unclear and incomplete information when merely relying on terminal fragment ions. Here, we show with spectra of synthetically generated histone peptides and proteins, how much information can be gained from including internal ions. By using long, 50 residues long sequences of commonly heavily modified H3.1 histone proteins, while knowing the content of the spectra, we are able to assess the value of our approach in a typical setting where the position of a modification on the amino-acid sequence is unknown. We will then assess the validity of the method in spectra from top-down MS.

In the process of annotating internal ions in long sequences, the number of theoretical fragments significantly surpasses the number of peaks observed in the experimental spectrum. This results in numerous mass overlaps among theoretical fragments, and therefore in competition between these fragments, as multiple theoretical fragments can match the same experimental peaks. In this competitive context, increasing the amount of considered theoretical fragments can reduce false positive fragment annotations. The rationale behind this is that higher cosine similarity between theoretical and experimental isotopic distribution denote more correct theoretical fragments. As the cosine similarity is used to filter annotations, this prioritizes spectral peaks annotation to correct fragments and limits annotation of incorrect fragments. Consequently, when comparing the annotation of fragments from various candidate proteoforms, it is more effective to annotate all theoretical fragments collectively in the spectrum. The graph-based annotation strategy described employs this method by allowing the input of multiple proteoforms and by integrating their theoretical fragments into a single graph. From a computational perspective, this strategy offers the advantage of eliminating redundant matching, thereby accelerating the annotation process.

It is important to emphasize that the inclusion of internal ions in data analysis necessitates careful evaluation and may not be useful in every context. Particularly, the use of internal ions for the identification of the primary amino acid sequence can make the search and scoring of candidate species much more time-consuming and prone to false-positives annotations. When annotating incorrect sequences where most theoretical fragments are not expected in the experimental spectrum, many internal fragments get annotated to peaks corresponding to actual terminal fragments. We illustrate that problem, and the extent of false-positive internal fragment annotation it leads to, by annotating a spectrum with a shuffled sequence of the expected sequence 6A. However, when annotating the spectrum with fragments from the correct sequence, experimental peaks tend to be assigned to the terminal fragments first, which reduces the number of internal fragments annotated and potentially the false-positive annotations 6A. This implies that internal fragments could be used in scenarios where the majority of the annotated peaks are correct and only a small subset of theoretical fragments requires comparison. Such circumstances arise notably in the localization of PTMs. In PTM localization problems, only a subset of fragment ions are significant for determining the modification site.

To evaluate if internal ions can boost the confidence in PTM localization, we modeled a situation where the position of a modification is unknown and two potential modification sites must be compared. To simulate a wrong modification, we introduced a mass shift in the searched peptidoform between near amino acids and compared the resulting intensities changes of site-determining internal and terminal ions with those from the correct proteoform (i.e. the proteoform without the mass shift)(Fig 7A). The comparison of fragment intensities done for each mass shift is illustrated in the fragmentation matrices of Fig. 6B and supplementary Fig. S7. The results of this test on a H3 spectrum are presented in Fig. 6B and show that a significant decrease in internal site-determining ions intensity is observable only when the correct original sequence is used for the annotation but not when the shuffled sequence is used. This mass-shift procedure was carried out on the entire sequence on middle-down and top-down spectra. The differences in intensity were investigated for several characteristic mass shifts corresponding to common modifications (Methylation, Acetylation, and Phosphorylation). The distance between amino acids (3, 5, and 7 residues apart) undergoing the mass shift was varied to test the sensitivity of the modification localization, as near modification sites present fewer site-determining ions, and therefore tend to be more difficult to distinguish.

**Figure 6:**
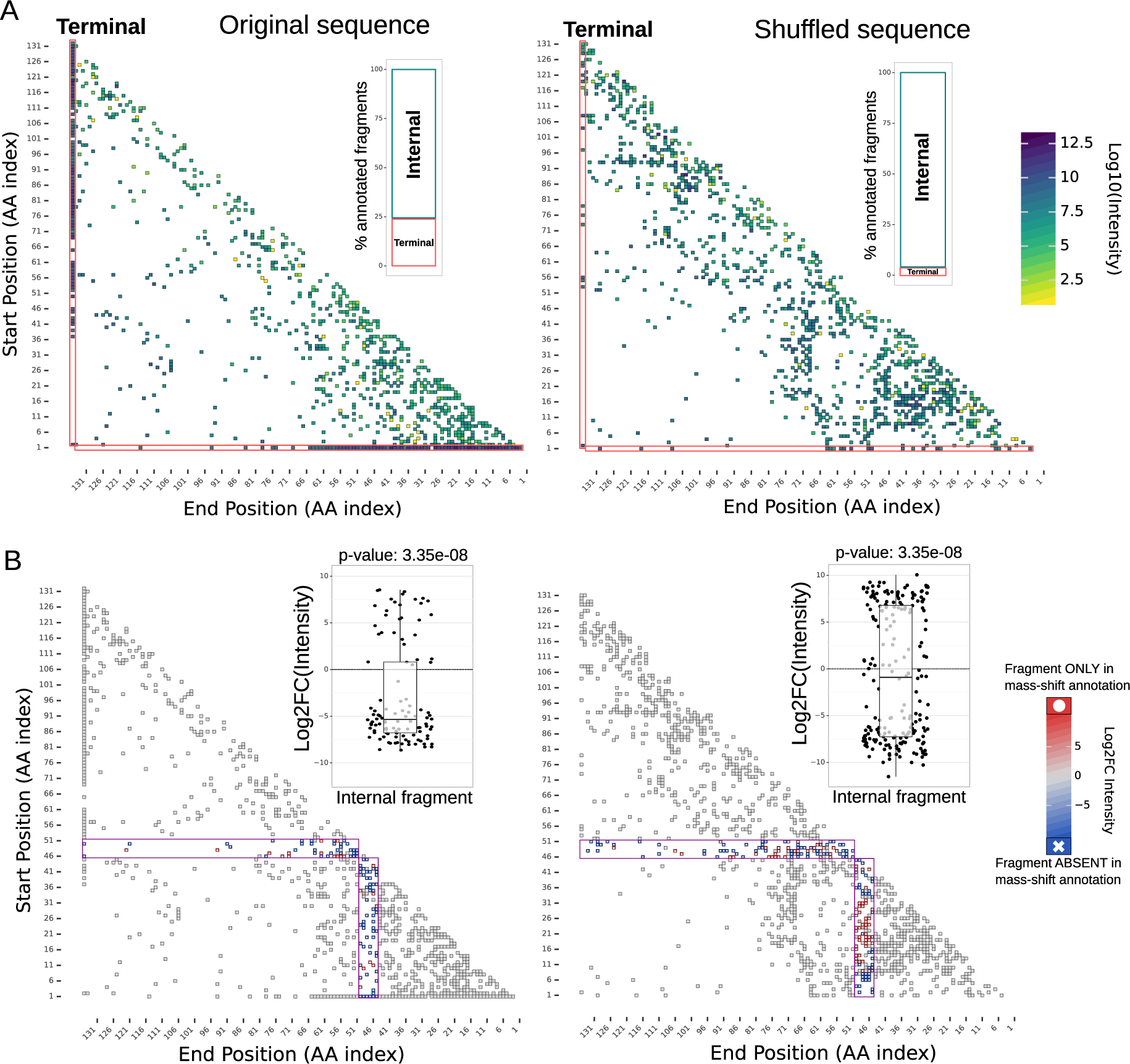
Significant change in internal ions intensities is observed when correct and incorrect fragments are competing. **A:** Annotated fragment intensity in histone H3 spectrum. The spectrum was annotated using the original and a shuffled sequence. The annotation using a shuffled sequence results in a much higher proportion of internal ions annotations (barplot). This is most likely due to the false-positive annotation of internal ions to peaks corresponding to terminal ions (red frames). **B:** Fragment intensity changes were measured after inducing a mass shift corresponding to phosphorylation (79 Da) between residues 45 and 51. Significant changes in internal fragment intensities can be observed only in the original sequence annotation (box plots).

**Figure 7:**
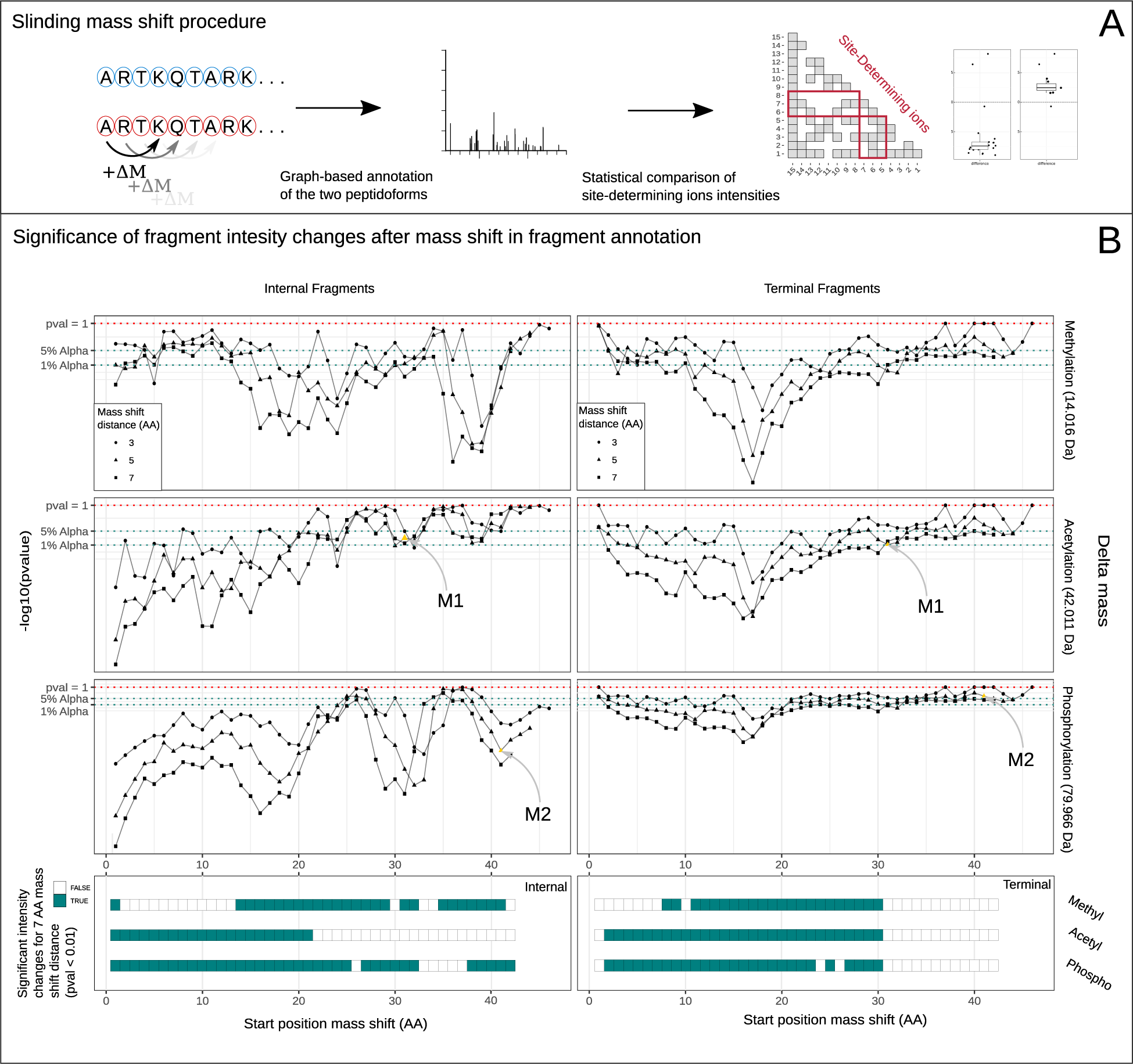
Internal ions provide relevant information for the localization of PTMs at different regions. **A:** The procedure referred to as “sliding mass shift” has been used to test whether site-determined ions presented a significant change in intensity when a mass shift was induced. This consisted in annotating both the expected peptidoform and the peptidoform where a mass shift (Delta mass) between two amino acids has been included. **B:** Significance level for different Delta masses (corresponding to Acetylation, Methylation, and Phosphorylation) and mass shift distances, for both internal and terminal ions are presented. **C:** Summary of the results for mass shift distance of 7 amino acids for the 3 delta masses where the mass shifts induced significant changes (p=0.01, fragment mass difference against 0 level, one-sided t-test) in the fragment intensities. (Dataset: Middle-down Histone H3 n-term tail, ETD 30ms). Example of intensity change fragmentation matrices for datapoint M1 and M2 are presented in Supp. Fig. S7.

The comparison of the annotated fragments between the original sequence and the sequence with a mass shift reveals significant changes in the intensities of site-determining fragments, for both internal and terminal fragment ions (Fig. 7B). Although the intensity of internal ions generally tends to be lower than that of terminal ions (usually about a 10-fold difference in middle-down data, ETD 30ms), the high number of site-determining internal ions confers an advantage over terminal fragments when statistically assessing changes in their intensity. The greater the distance between amino acids undergoing the mass shift results in more site-determining ions and, consequently, more significant changes in intensity. Figure 7B illustrates that the considered mass shift significantly influences the region of the sequence where substantial changes in internal ions are observed. An important observation is the complementary nature of information from internal ions to that of terminal ions. The region of the sequence with significant changes in intensity is summarized in the lower part of Fig7B. Some sequence regions of the sequence might show no change in terminal ion intensities but exhibit significant changes in internal ion intensities. These are most likely regions where internal ions provide relevant additional information. As a control, the same procedure was applied using a shuffled sequence of H3 n-terminal tail (Suppl. Fig. S8).

We will now investigate whether internal fragments can be beneficial for PTM localization in top-down MS. When applying the same procedure to top-down spectra of synthetic histone H3 (Fig. S9), we observe less significant changes in internal ions around the PTM. This can be attributed to the two following reasons. First, the spectra’s complexity is higher due to double to triple the sequence lengths, leading to increased fragment overlap and a higher likelihood of annotating noise peaks as shown in Fig. 6. Secondly, the fragmentation parameters employed for generating the MS2 scans in these experiments were not optimized for internal ion generation (ETD 30ms for the spectra in Fig. 7, compared to 15ms for Fig. S9), resulting in supposedly fewer generated internal ions. Despite their potentially lower abundance, as displayed in Fig. S9 internal ions still provide valuable insights into PTM localization for the N-terminus of the sequence, where terminal ions show less pronounced changes in annotated site-determining ions.

Summarizing the outcomes of the sliding mass shift procedure illustrated for both middle-down and top-down spectra (Table 1), it becomes evident that the previously highlighted “competition” among theoretical fragments has to be considered. In the annotation method where two peptidoforms are simultaneously annotated, there is a notable 15% increase in the number of significant intensity difference in the internal ions, observed in both deconvoluted and centroided spectra. Concerning deconvolution, valuable information appears to be compromised, as approximately 30% fewer tests are significant in deconvoluted spectra but register as significant in centroided ones. The control using a shuffled sequence indicates that it is still possible to obtain significant results, this can partly be the result of the randomized sequence still containing a combination of amino-acids fragments that are found in the original sequence. Finally results in the top-down dataset display less significant change in site-determining fragment intensities; the proportion of internal fragments might not be optimal due to the low ETD reaction time used to generate these spectra.

**Table 1:** Summary of sequence regions displaying significant change in fragment intensities after induced mass. The table displays the percentage of significant differences, at 1% alpha level from t-tests, for each comparison of fragment intensities performed in the sliding mass shift experiment. This was performed the following two annotation methods; Competitive: where both peptidoform theoretical fragment lists are annotated simultaneously in the spectra, Distinct: two independent annotations are performed. The comparison was performed from both deconvoluted spectra and non-deconvoluted spectra (denoted “centroided” here). The shuffled sequence of the H3 N-terminal tail is used here as a double control. Results for top-down spectra (H4, H3, and Filgrastim) are only shown for the best-performing method.

| Protein/Peptide | Sequence:<br>Original/Shuffled | Annotation method | Pre-processing | p.val. from | Percentage<br>significant tests |
| --- | --- | --- | --- | --- | --- |
| H3 N-tail | **Original** | **Competitive** | **Centroided** | **Internal** | 57.1 |
| H3 N-tail | **Original** | **Competitive** | **Centroided** | Terminal | 40.2 |
| H3 N-tail | **Original** | **Competitive** | Deconvoluted | **Internal** | 16.4 |
| H3 N-tail | **Original** | **Competitive** | Deconvoluted | Terminal | 39.6 |
| H3 N-tail | **Original** | Distinct | **Centroided** | **Internal** | 31.9 |
| H3 N-tail | **Original** | Distinct | **Centroided** | Terminal | 36.4 |
| H3 N-tail | **Original** | Distinct | Deconvoluted | **Internal** | 1.8 |
| H3 N-tail | **Original** | Distinct | Deconvoluted | Terminal | 29.5 |
| H3 N-tail | Shuffled | **Competitive** | **Centroided** | **Internal** | 14.1 |
| H3 N-tail | Shuffled | **Competitive** | **Centroided** | Terminal | 2.5 |
| H4 | **Original** | **Competitive** | **Centroided** | **Internal** | 15.3 |
| H4 | **Original** | **Competitive** | **Centroided** | Terminal | 40.6 |
| H3 | **Original** | **Competitive** | **Centroided** | **Internal** | 17.5 |
| H3 | **Original** | **Competitive** | **Centroided** | Terminal | 25.2 |
| Filgrastim | **Original** | **Competitive** | **Centroided** | **Internal** | 1.9 |
| Filgrastim | **Original** | **Competitive** | **Centroided** | Terminal | 21.1 |

## 5 Discussion

Internal fragments, despite their lower intensity compared to terminal ions, can constitute a significant portion of spectral information in middle-down and top-down MS spectra. Their large number offers a potential statistical advantage for deep proteoform characterization. We showed that utilizing internal ions requires careful consideration of their potential for adding false positive information. It is important to emphasize that annotating internal ions without stringent filtering will almost always enhance sequence coverage from spectra of long peptides or entire proteins by adding many false annotations of spectral peaks when considering all internal ions.

On top of false positives, it can also be hard to annotate the correct fragment ions. While advancements in mass spectrometer resolution are beneficial, this does not resolve the problem of overlapping fragments. Higher resolution would improve the distinction of near-isobaric fragments, and better signal-to-noise ratios would limit false-positive annotation to noise. However, we show here that a major challenge resides in the combinatorial nature of the problem, where fragment overlaps often are the result of exact mass coincidences that cannot be resolved by narrower tolerance windows.

To address these issues and improve the usability of internal ions, we present a set of methods to represent, visualize and compare fragmentation spectra with internal ions. We introduced a strategy that leverages the complementarity between terminal and internal ions. This approach can be employed to tune fragmentation parameters to maximize the generation of secondary internal ions without compromising terminal fragment information. This will lead to higher coverage and less likely falsely identified internal fragment ions.

The developed graph-based annotation method allows to pinpoint key aspects that can be considered to improve internal fragment annotation and usage. The fragmentation graph addresses the considerable information loss in the deconvolution step. The annotation of non-deconvoluted spectra through the representation of all possible theoretical fragments opens the door for more accurate scoring methods for assessing their existence. Using non-deconvoluted spectra allows to score fragment annotation by comparing expected and measured isotopic patterns for efficient filtering strategies. Most importantly, the rate of false-positive annotations can be reduced by direct comparison of different proteoforms on the same graph. With such a graph annotation, the proteoforms “compete” for being more representative and more explained. Using this approach for annotation, we demonstrate use cases where internal ions provide valuable insights for PTM localization. With the analysis of intensity changes in specific site-determining ions across candidate proteoforms, it is possible to statistically ascertain the relevance of internal fragment information. Essentially, this approach provides a statistical framework for helping in the localization of PTM using internal ions. We show its superiority for histone N-terminal tails in middle-down data and show its high potential for longer peptides and full proteins in top-down proteomics data, particularly after optimizing the fragmentation parameters. Overall, the methods described in this work can be used to enhance proteoform characterization and clarify PTM assignments on long species identification, and possibly lay the groundwork for integrating internal ions into the scoring or rescoring of spectra identifications.

## 6 Acknowledgments

We would like to thank Louise Marie Buur, Micha Birklbauer, Zoltan Udvardy, and Caroline Lennartsson who took part in the project “Making sense of internal fragment ions” of the EuBIC-MS developers meeting edition 2023. This project has received funding from the European Union Horizon 2020 research and innovation program under the Marie Sk-lodowska-Curie grant agreement No 956148. Proteomics and mass spectrometry research at SDU is supported by generous grants to the VILLUM Center for Bioanalytical Sciences (VILLUM Foundation grant no. 7292 to ONJ), PRO-MS: Danish National Mass Spectrometry Platform for Functional Proteomics (grant no. 5072-00007B to ONJ) and INTEGRA (Novo Nordisk Foundation, grant no. NNF20OC0061575 to O.N.J).

## 7 Data Availability

The mass spectrometry proteomics data have been deposited to the ProteomeXchange Consortium (http://proteomecentral.proteomexchange.org) via the PRIDE repository. The histone H3 N-terminal tail dataset is deposited under the dataset identifier PXD050421. The intact histones dataset is deposited under the identifier PXD050615. The data are available in Thermofisher raw files and mzML format. The source code of the fragmentation graph procedure as well as a vignette describing how to use the tool is available at https://github.com/arthur-grimaud/fragmentation-graph/ (DOI: 10.5281/zenodo.10785142). Additional analysis of the fragmentation graph output and analysis of theoretical fragments has been conducted in R. The source code for these has been deposited in another GitHub repository https://github.com/arthur-grimaud/fragmentation-graph manuscript archive.

## Abbreviations

PTM: post-translational modification
MS: mass spectrometry
MS2: Tandem mass spectra
ETD: electron-transfer dissociation

