## Supplementary Material for "How to deal with internal fragment ions?"

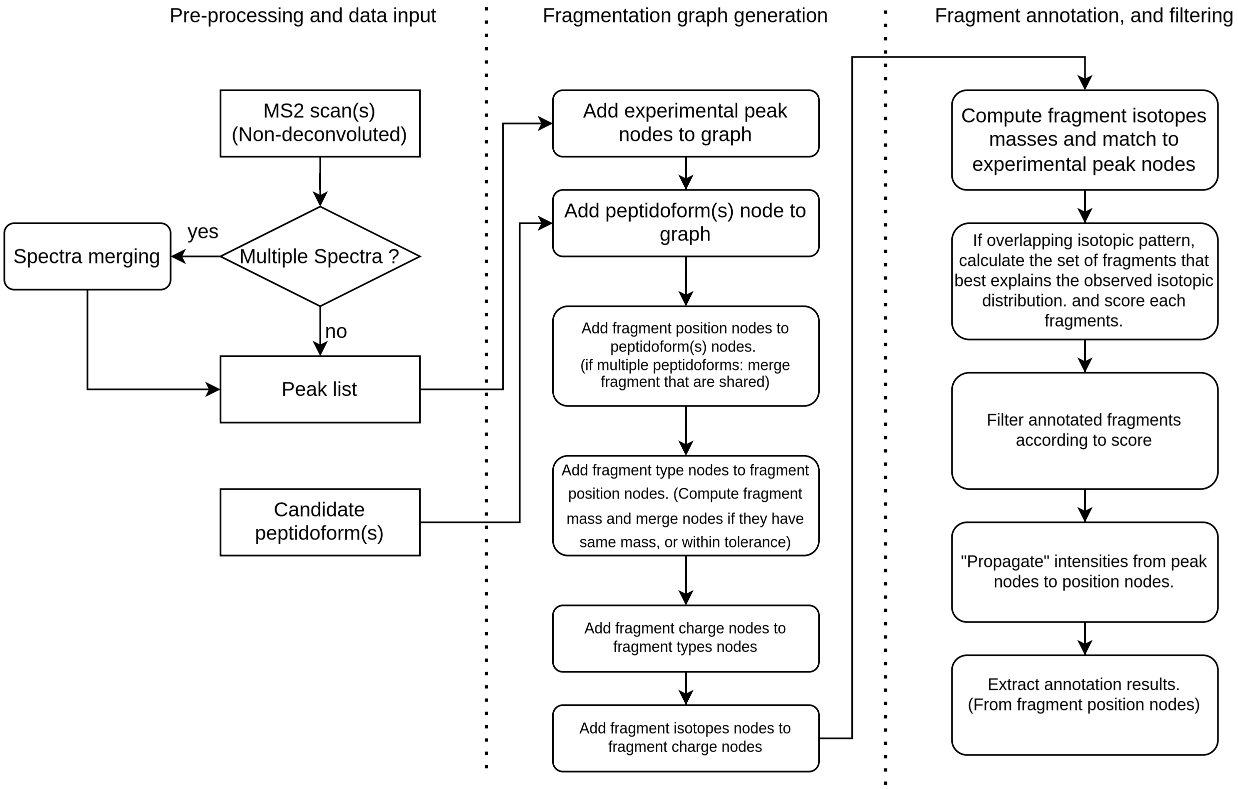


Figure S1: **Diagram of the fragmentation graph annotation procedure.** The diagram describes the main steps from the pre-processing of spectra to the generation of the fragmentation graph and annotation results.

| **Ion cap** | **Formula** | **Delta Mass (Da)** |
|---|---|---|
| a | H-2O-1C-1O-1 | -46.00548 |
| b | H-2O-1 | -18.01056 |
| x | H-2O-1CO2 | 25.97926 |
| y | | 0.00000 |
| c-1 | H-2O-1NH3H-1 | -1.99184 |
| c | H-2O-1NH3 | -0.98402 |
| cdot | H-2O-1NH3H1 | 0.02381 |
| c+1 | H-2O-1NH5 | 1.03163 |
| zdot | H-2O-1N-1OH-1 | -17.02655 |
| z+1 | H-2O-1N-1O | -16.01872 |
| z+2 | H-2O-1N-1OH | -15.01090 |
| z+3 | H-2O-1N-1OH2 | -14.00307 |

Table S1: **Ion caps delta mass used in fragment mass calculation** The table presents the formula (in *pyteomics.mass* package format) used in the fragmentation graph to compute the delta mass of the different fragment ions. The delta masses are given in Dalton (rounded value at the 5th decimal). For example the ion cap of an "a" c-terminal fragment results in a mass loss of 46.00548 Da equivalent to a loss of one H2O and one CO group.

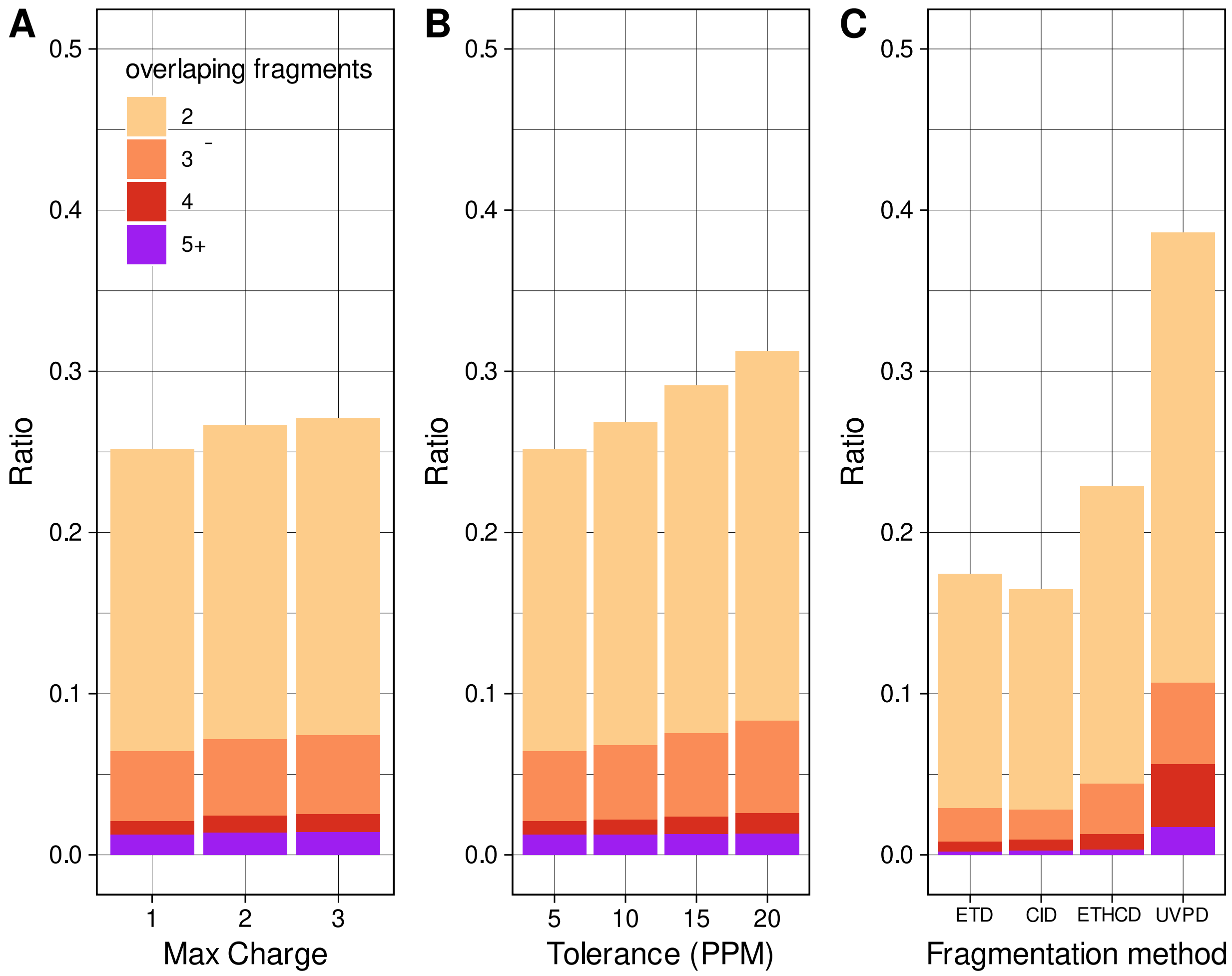


Figure S2: **Effect of various parameters on the rate of fragment overlap.** The average ratio of overlapping fragments in theoretical spectra of peptides of length 20 was computed for different max charge states (A), Tolerance (B), and Fragmentation Method ( C). The fragment types included for each fragmentation method are given in Supplementary 7. If not specified otherwise in the figure the following settings were used: max charge: 1, tolerance: 5PPM, fragmentation method: ETD.

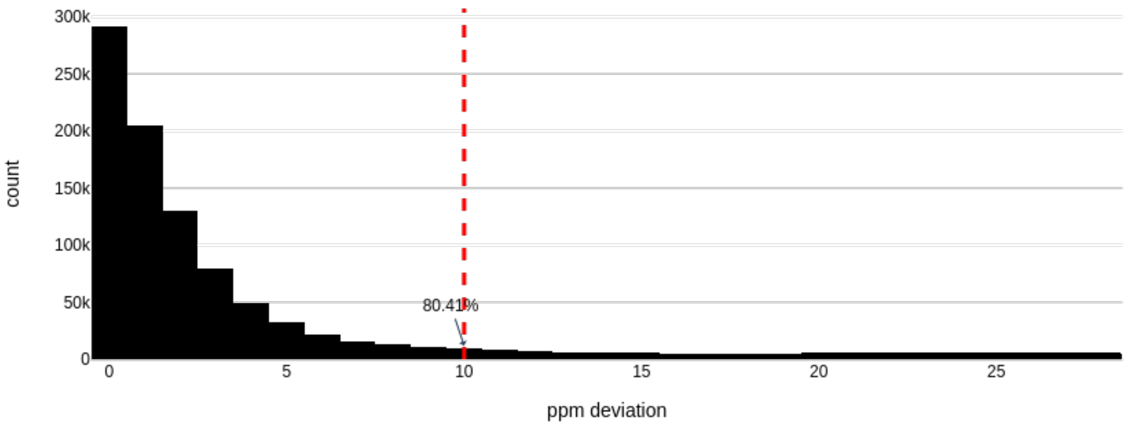


Figure S3: **Determination of MSMS tolerance for spectra merging.** For each peak of the spectra to be merged the distance to the closest peak in the other spectrum is computed and represented here in a histogram. This is used to determine the optimal m/z tolerance for spectra merging

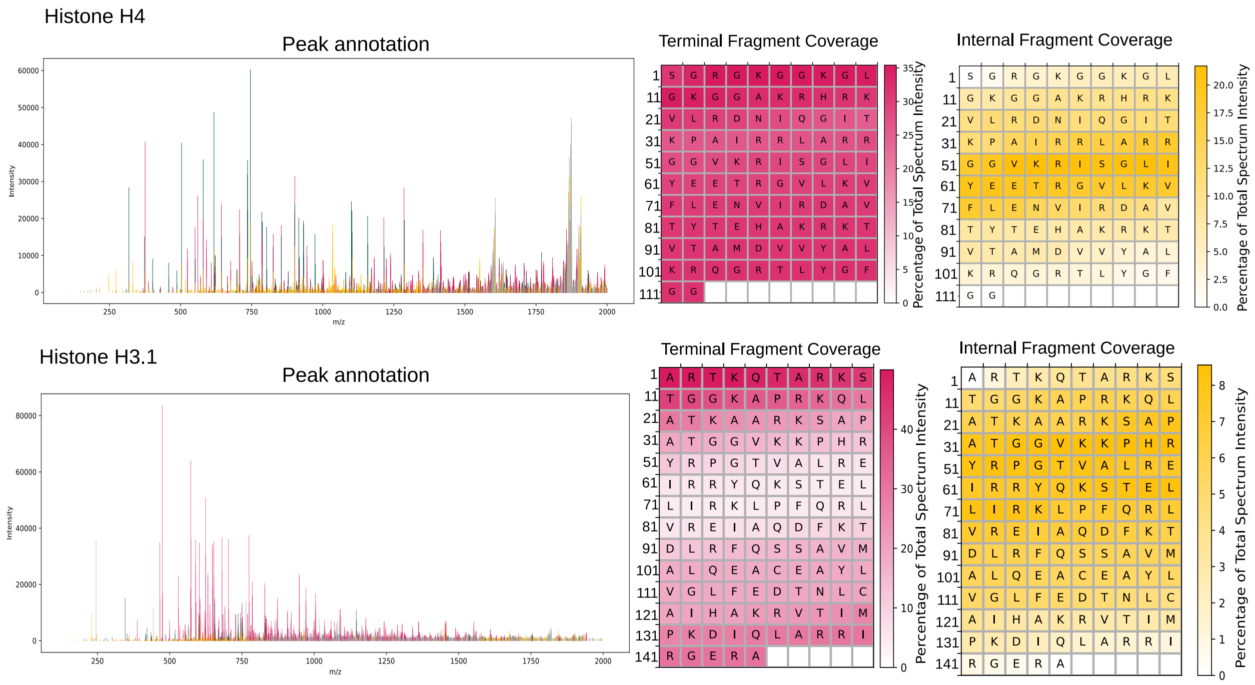


Figure S4: **Annotation of internal fragments enhances sequence coverage in entire protein spectra.** Spectra have been annotated using the fragmentation graph procedure in the consensus spectrum of Histone H4 (**TOP**, ETD 15ms, 5 merged ms2 scans) and Histone H3.1 (**BOTTOM**, ETD 15ms, 5 merged ms2 scans). Sequence coverage is represented as the percentage of intensity in the MS2 scan covering each residue, internal and terminal fragment coverages are presented separately.

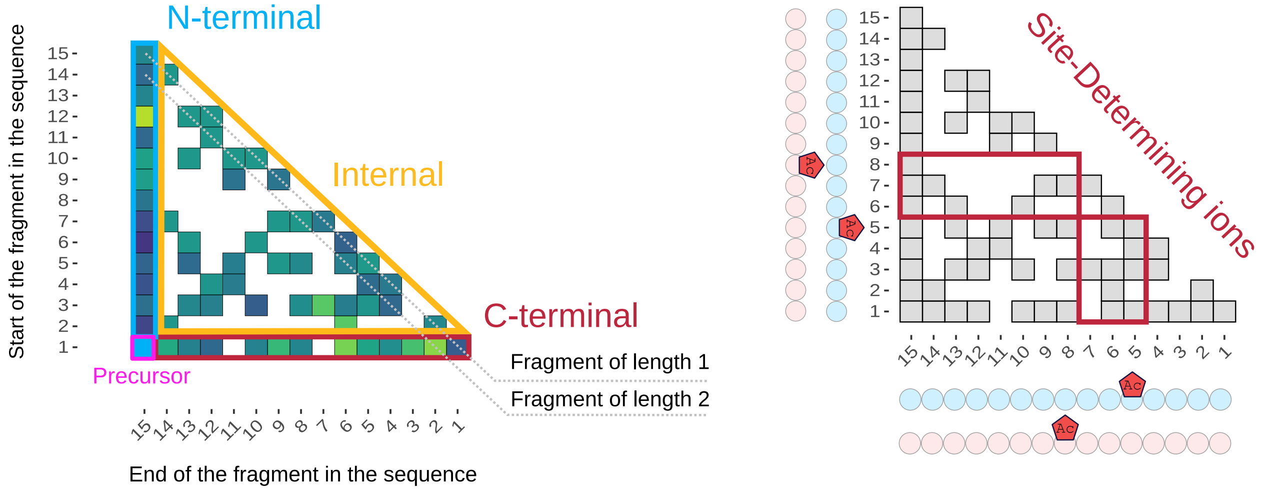


Figure S5: **How to read a fragmentation matrix. A:** Fragmentation matrix provides an easy way to visualize all fragments annotated. In that representation, the y-axis indicates the index of the starting amino acid of the fragment in the sequence and the x-axis the last amino acid of the fragment in such a way that the first column displays the N terminal fragment intensities and the Bottom row the C terminal fragment intensities. All the other cells represent internal ions and fragments of the same length are present on the same diagonal (dashed line). **B:** That representation is also convenient when comparing two annotations with different modification positions for instance. The site-determining ions can be easily visualized (red boxes, **B**).

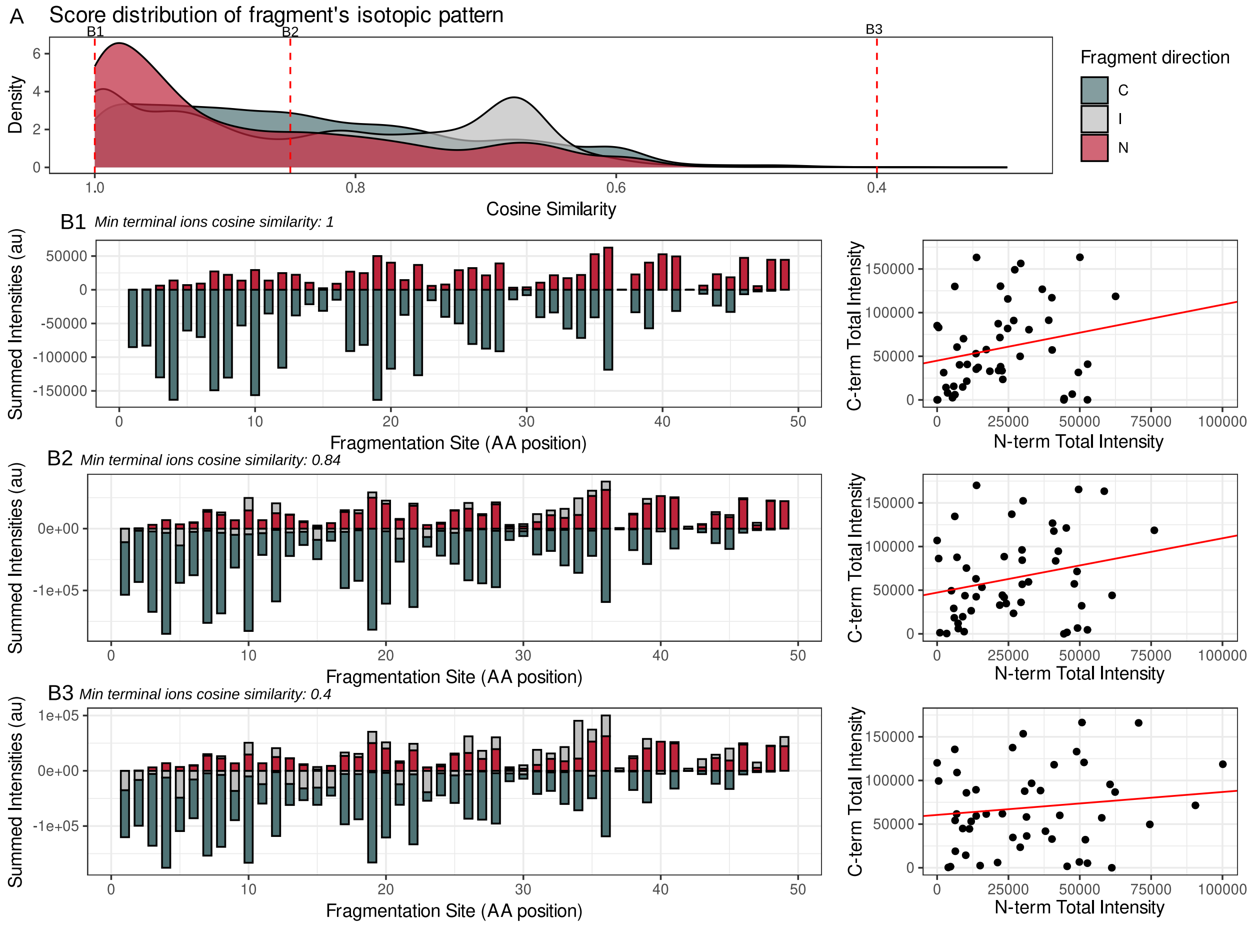


Figure S6: **Annotation of internal in MS2 scan from weak fragmentation does not improve the correlation between N and C terminal intensity of fragmentation site.** In opposition to the result shown for ETD 40ms, the annotation of internal fragments in peak-picked spectra of low fragmentation intensity (N=45, ETD reaction time 10; supplementary activation 10) does not improve the correlation between N and C term intensities. In such MS2 scans the occurrence of secondary fragmentation events yielding internal ions is probably very low, and therefore these internal fragments probably represent a non-significant fraction of the MS2 scan intensities making their annotation not relevant.

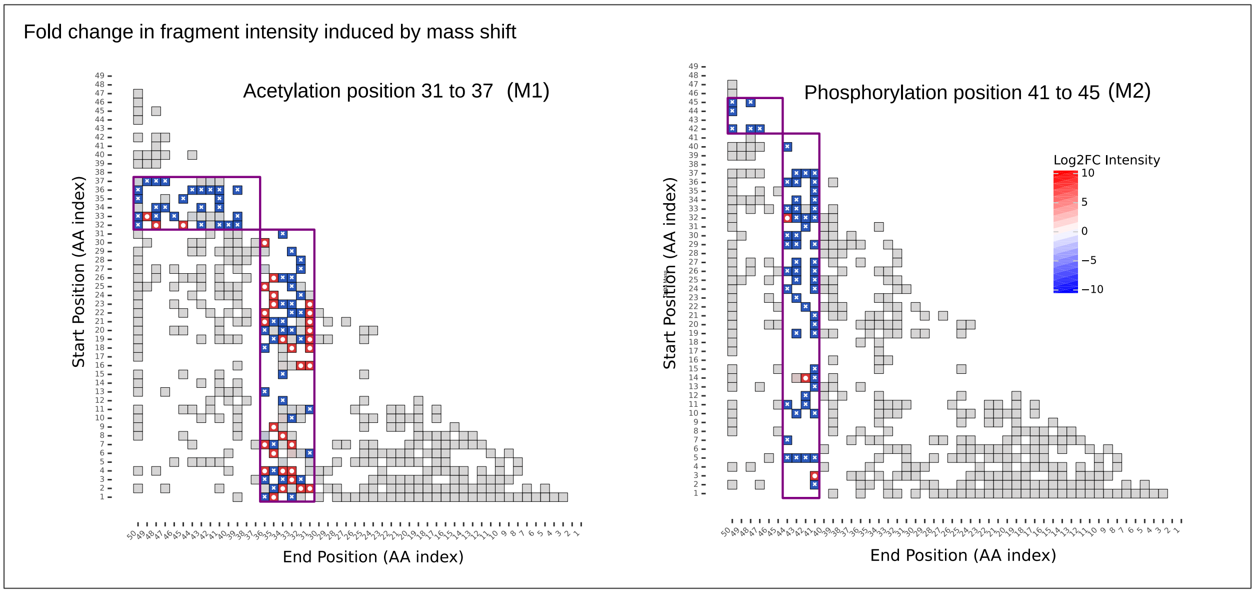


Figure S7: **Fragment intensity change matrix.** Example of two fragment intensity matrix used in the “sliding mass shift” procedure described in figure 6. Blue display decreases in intensity or a loss of the annotation for fragments where the mass shift has been induced. Red display fragments whose intensity increases after the mass shift is induced.

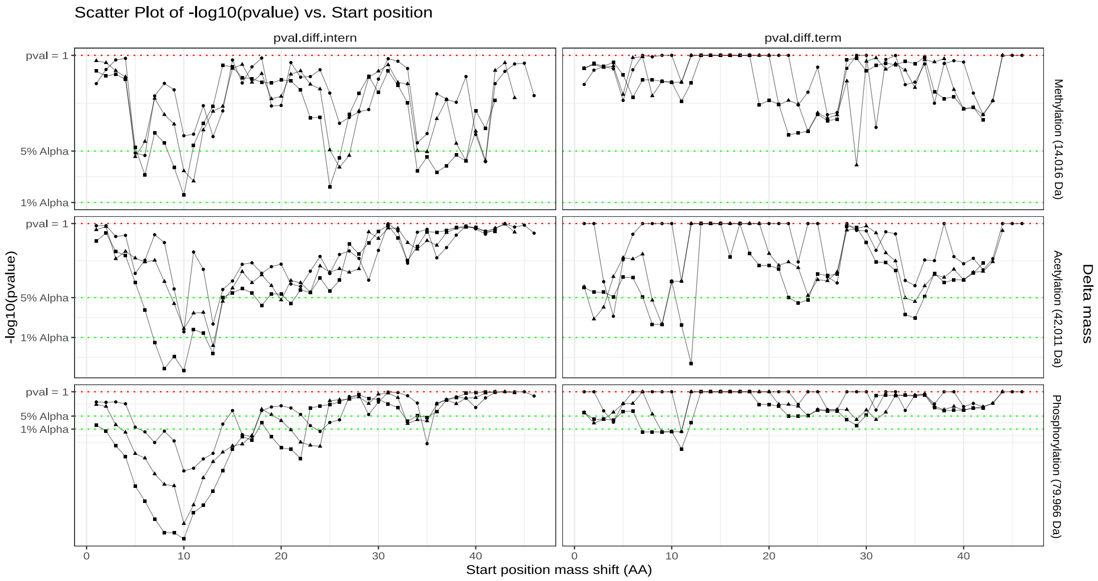


Figure S8: **Sliding mass shift test in H3 N-tail spectra using shuffled sequence as a decoy.** Mass shift procedure was applied to middle down Histone H3 n terminal tail spectra, theoretical fragment where generated from the shuffled original sequence

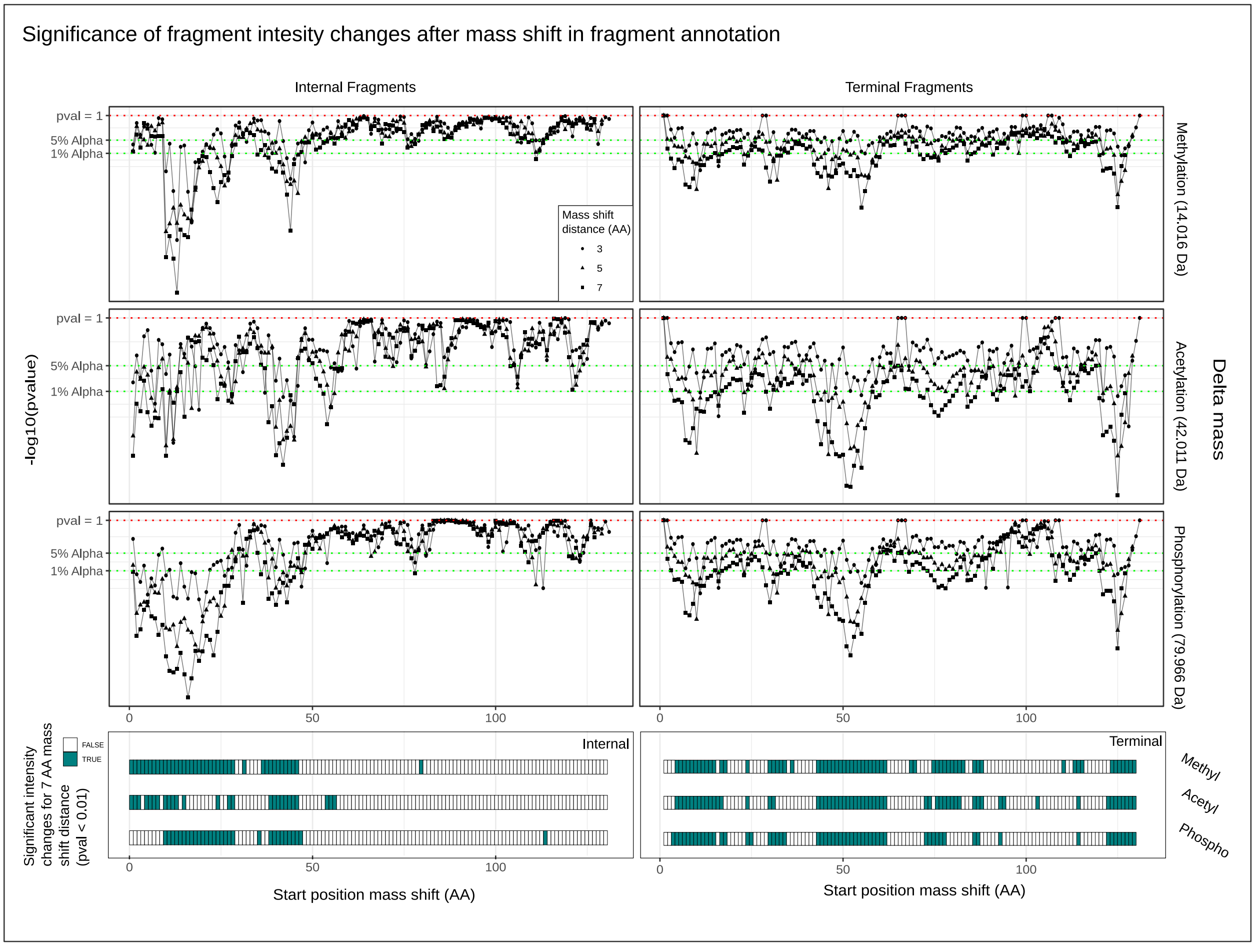


Figure S9: **Comparison of internal and terminal ions information for the localization of PTMs in Top-down data using the "sliding mass shift procedure".** Dataset: Top-down: Histone H3, ETD 10ms, 15 spectra merged in consensus spectra

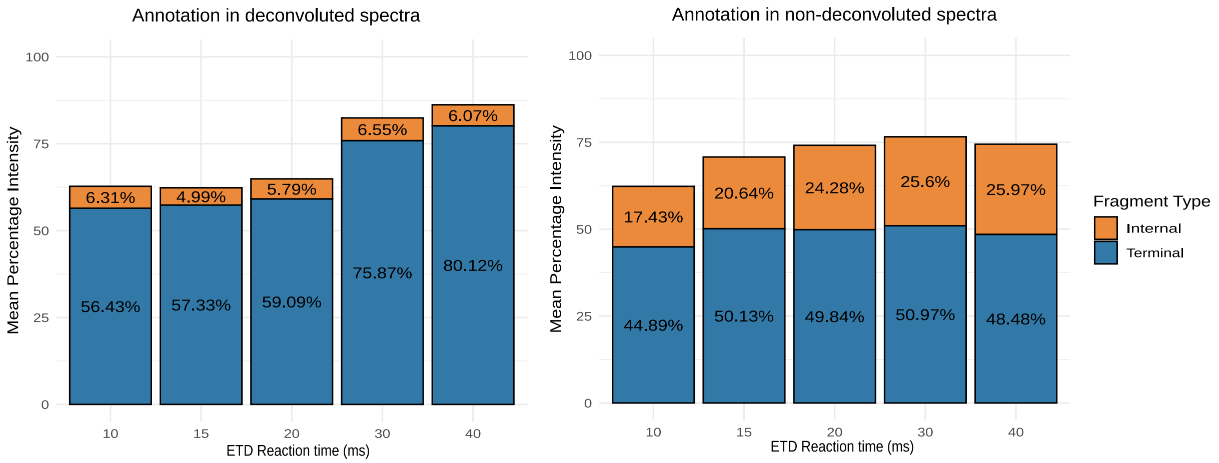


Figure S10: **Deconvolution reduces available information on internal ions**Average intensity assigned to internal and terminal for deconvoluted and non-deconvoluted spectra (dataset:middle-down histone h3 N terminal tail,n=45)
